# PyOrthoANI, PyFastANI, and Pyskani: a suite of Python libraries for computation of average nucleotide identity

**DOI:** 10.1101/2025.02.13.638148

**Authors:** Martin Larralde, Georg Zeller, Laura M. Carroll

**Author notes:** Corresponding authors: Martin Larralde,; Laura M. Carroll.

## Abstract

**Summary:** The average nucleotide identity (ANI) metric has become the gold standard for prokaryotic species delineation in the genomics era. The most popular ANI algorithms are available as command-line tools and/or web applications, making it inconvenient or impossible to incorporate them into bioinformatic workflows, which utilize the popular Python programming language. Here, we present PyOrthoANI, PyFastANI, and Pyskani, Python libraries for three popular ANI computation methods. ANI values produced by PyOrthoANI, PyFastANI, and Pyskani are virtually identical to those produced by OrthoANI, FastANI, and skani, respectively. All three libraries integrate seamlessly with BioPython, making it easy and convenient to use, compare, and benchmark popular ANI algorithms within Python-based workflows.

**Availability and Implementation:** Source code is open-source and available via GitHub (PyOrthoANI, https://github.com/althonos/orthoani; PyFastANI, https://github.com/althonos/pyfastani; Pyskani, https://github.com/althonos/pyskani).

**Supplementary Information:** Supplementary data are available on *bioRxiv*.

## Introduction

The average nucleotide identity (ANI) metric of genomic similarity is arguably the most popular method for prokaryotic species delineation in the genomics era (Jain *et al*., 2018; Richter and Rosselló-Móra, 2009). The calculation of ANI values shared between two genomes is a crucial step in many bioinformatic pipelines, including popular methods/workflows for prokaryotic species identification (Parks *et al*., 2018; Chaumeil *et al*., 2019), within-species lineage/strain delineation (Rodriguez-R *et al*., 2024; Raghuram *et al*., 2024), and general prokaryotic (meta)genomic data analysis (Olm *et al*., 2017; Petit and Read, 2020).

While numerous ANI algorithm implementations have been developed, nucleotide BLAST-based ANI (ANIb) algorithms are considered to be the gold standard (Jain *et al*., 2018; Shaw and Yu, 2023). ANIb algorithms are accurate in the sense that they share a strong correlation with experimentally determined DNA-DNA hybridization (DDH) values (Konstantinidis and Tiedje, 2005; Goris *et al*., 2007; Lee *et al*., 2016; Richter and Rosselló-Móra, 2009). However, due to the high time complexity of BLAST and similar alignment-based algorithms, ANIb algorithms are notoriously slow (Jain *et al*., 2018) and thus most appropriate for users with smaller datasets (e.g., up to ≈10^3^ genomes/10^6^ pairwise comparisons), who prioritize accuracy over speed.

To overcome the computational limitations of ANIb, alignment-free ANI algorithms have been developed, most notably FastANI (Jain *et al*., 2018) and skani (Shaw and Yu, 2023). Both FastANI and skani forgo some accuracy in favor of speed (i.e., they produce ANI values, which correlate with, but are not necessarily equivalent to, ANIb), and as such, they can readily scale to massive genomic datasets (e.g., ≥10^4^ genomes/10^8^ pairwise comparisons) (Jain *et al*., 2018; Shaw and Yu, 2023). However, identifying the optimal alignment-free ANI algorithm for a given dataset is not always straightforward. FastANI is ≥50x faster than ANIb methods and is more accurate than skani on reference-quality genomes (Shaw and Yu, 2023; Jain *et al*., 2018). skani, on the other hand, is >20x faster than FastANI and is more accurate on fragmented, incomplete metagenome-assembled genomes (MAGs) (Shaw and Yu, 2023). Thus, in addition to considering dataset size and algorithm speed-accuracy tradeoff, users may want to consider dataset composition (e.g., isolate genomes versus MAGs) and quality when selecting the optimal ANI algorithm for their dataset.

Regardless of whether they prioritize accuracy or speed, the most popular ANI algorithms/methods (e.g., FastANI, skani, ANI by Orthology [OrthoANI], JSpeciesWS, PyANI) are available as command-line tools and/or web applications (Jain *et al*., 2018; Shaw and Yu, 2023; Lee *et al*., 2016; Richter *et al*., 2016; Pritchard *et al*., 2015). This makes it inconvenient–and sometimes, impossible–for bioinformaticians to incorporate ANI algorithms into bioinformatic workflows, which utilize the popular Python programming language (Van Rossum and Drake, 2009).

Here, we present a suite of Python libraries for popular ANI algorithms, specifically: (i) PyOrthoANI, a Python-based implementation of the OrthoANI algorithm (a highly accurate ANIb method) (Lee *et al*., 2016); (ii) PyFastANI and (iii) Pyskani, Python bindings for the FastANI and skani algorithms, respectively (fast, alignment-free methods) (Jain *et al*., 2018; Shaw and Yu, 2023). Each Python library integrates seamlessly with BioPython (Cock *et al*., 2009), making it simple and convenient to perform ANI computations within Python-based bioinformatic workflows, software programs, and notebooks (e.g., Jupyter) (Kluyver *et al*., 2016). By providing a unified Python interface, our suite allows users to easily swap out different ANI algorithms, making it simple and convenient to test, compare, and benchmark methods.

## Implementation

The PyOrthoANI algorithm (https://github.com/althonos/orthoani) was implemented in the same manner as the original OrthoANI Java implementation (Lee *et al*., 2016).

Briefly, to calculate ANI values between a query and reference genome, both genomes are partitioned into 1,020 bp-long fragments. Fragments that are <1,020 bp in length and/or contain >80% ambiguous (N) nucleotides are discarded. Nucleotide BLAST (blastn) (Camacho *et al*., 2009) values are then calculated between the set of query and reference genome fragments using the following blastn parameters (all other parameters are set to their respective defaults): -evalue 1e-15, -xdrop_gap 150, -dust no, -penalty -1, -reward 1, -num_alignments 1, -outfmt ‘6 qseqid sseqid length pident’.

The resulting fragments are considered to be orthologous if they produce reciprocal best hits, which cover at least 35% of the total length of the fragment. Final ANI values are calculated by averaging the nucleotide identity values for all reciprocal blastn hits.

For PyFastANI (https://github.com/althonos/pyfastani), the original FastANI code (written in C++) (Jain *et al*., 2018) was wrapped into a Python extension module using the Cython language (v3.0) (Behnel *et al*., 2011). While PyFastANI uses the original FastANI code for hashing and core-genome identity computations, we reimplemented the sketching to support passing plain Python strings as input sequences. In addition, we implemented serialization/deserialization support to allow querying a reference database several times. To speed up the querying of individual sequences, we parallelized the fragment sketching step using Python thread pools and re-entrant code.

For Pyskani (https://github.com/althonos/pyskani), the original skani code (written in Rust) (Shaw and Yu, 2023) was wrapped into a Python extension module using the PyO3 library (v0.22.5; https://pyo3.rs) for bindings generation. To accelerate querying, we implemented a more generic strategy for the storage of reference markers, allowing to either load the markers from a file iteratively (as in the original skani), or pre-loading them in memory to reduce I/O costs for successive querying.

## Results

Using each of the five data sets used to validate and benchmark FastANI (*n* = 14,952 total [meta]genomes) (Jain *et al*., 2018), we compared ANI values produced by PyOrthoANI, PyFastANI, and Pyskani to those produced by OrthoANI, FastANI, and skani, respectively. We additionally benchmarked the speed of all six methods on each genome individually using 1, 8, and/or 16 CPUs in triplicate (*n* = 717,651 total ANI computations; Figure 1, Supplementary Figures S1-S7, Supplementary Tables S1-S7, Supplementary Text).

**Figure 1.**
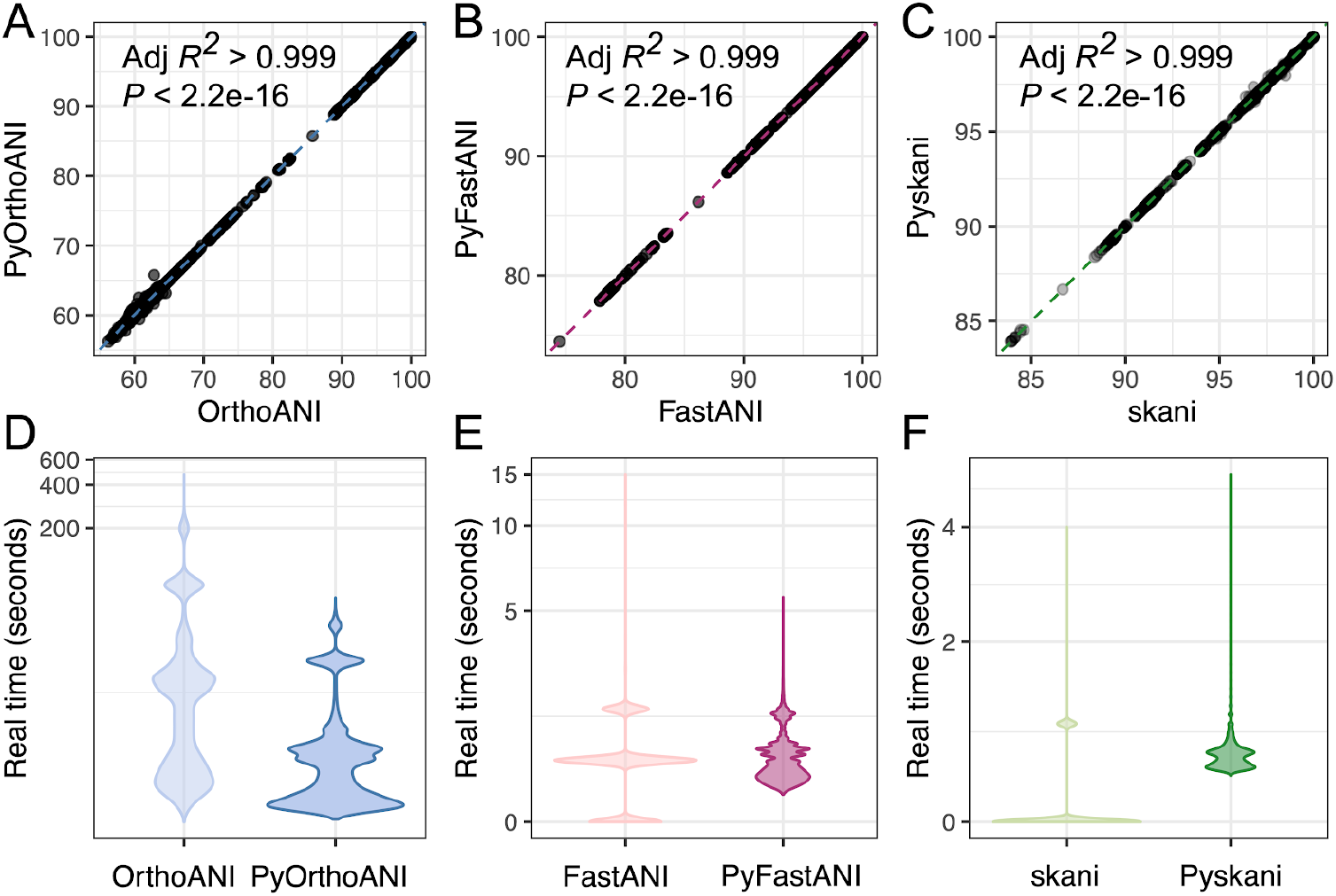
(A-C) Correlation between average nucleotide identity (ANI) values produced by (A) PyOrthoANI, (B) PyFastANI, and (C) Pyskani (Y-axes) with ANI values produced by OrthoANI, FastANI, and skani, respectively (X-axes) for genomes used in the FastANI validation/benchmarking data sets (black dots; Supplementary Tables S1-S6). Dashed lines denote the best-fitting linear model for each method pair, with adjusted *R*^2^ and *P*-values reported in the upper left corner of each subplot. Pyskani values were multiplied by 100. (D-F) Per-genome real (wall clock) time in seconds (Y-axes, log-scale) for (D) OrthoANI/PyOrthoANI, (E) FastANI/PyFastANI, and (F) skani/Pyskani (X-axes), using 1, 8, and/or 16 CPUs on the FastANI validation/benchmarking data sets (violin plots; Supplementary Table S7). For fairness, PyFastANI and Pyskani times include the time it took to load Python modules and parse genomes using BioPython (performed for every genome/computation). For extended versions of this figure, see Supplementary Figures S1-S7. Raw data used to construct all plots is available in Supplementary Tables S1-S7.

ANI values calculated by PyOrthoANI, PyFastANI, and Pyskani were virtually identical to those produced by OrthoANI, FastANI, and skani, respectively (adjusted *R*^2^ > 0.999 and *P* < 2.2e-16 for all methods; Figure 1a-c). Compared to OrthoANI, PyOrthoANI was, on average, 3x faster per genome (Figure 1d). PyFastANI and Pyskani performed similarly to FastANI and skani, respectively, even when Python module load times and genome parsing (via BioPython) were included in the PyFastANI/Pyskani runtime; however, differences in FastANI/PyFastANI and skani/Pyskani runtime and memory usage varied by dataset (Figure 1ef, Supplementary Figures S1-S7).

Overall, PyOrthoANI, PyFastANI, and Pyskani enable users to perform ANI computations within Python-based software, workflows, and notebooks. Because each Python library integrates with BioPython and is easily interchangeable, we anticipate that our Python suite will be particularly useful for comparing/benchmarking ANI algorithms, and for developers/users who frequently encounter highly heterogeneous datasets (e.g., genomic datasets varying in size, quality, and isolate/MAG composition) which require flexibility in ANI computation algorithms.

## Acknowledgements

This research was conducted using the resources of High Performance Computing Center North (HPC2N; Umeå University, Umeå, Sweden).

## Data availability

PyOrthoANI, PyFastANI, and Pyskani are available: (i) as part of the Python Package Index (PyPI) repository under the open-source MIT license at https://pypi.org/project/orthoani/, https://pypi.org/project/pyfastani/, and https://pypi.org/project/pyskani/, respectively; (ii) via GitHub (source code) at https://github.com/althonos/orthoani, https://github.com/althonos/pyfastani, and https://github.com/althonos/pyskani, respectively; (iii) as Singularity containers (used for benchmarking) at https://cloud.sylabs.io/library/lmc297/pyorthoani/pyorthoani, https://cloud.sylabs.io/library/lmc297/pyfastani/pyfastani, and https://cloud.sylabs.io/library/lmc297/pyskani/pyskani, respectively.

