## Supplementary Text for "PyOrthoANI, PyFastANI, and Pyskani: a suite of Python libraries for computation of average nucleotide identity"

for

by

Martin Larralde, Georg Zeller, and Laura M. Carroll

### Supplementary Methods

**Acquisition of (meta)genomic data for benchmarking.** Each of the five (meta)genomic data sets used to validate and benchmark FastANI (Jain *et al.*, 2018) were downloaded (*n* = 14,952 total [meta]genomes; http://enve-omics.ce.gatech.edu/data/fastani): (i) Dataset 1 (D1; NCBI RefSeq), with 1,662 genomes (http://enve-omics.ce.gatech.edu/data/public_fastani/D1.tar.gz); (ii) Dataset 2 (D2; *Bacillus cereus*), with 571 genomes (http://enve-omics.ce.gatech.edu/data/public_fastani/D2.tar.gz); (iii) Dataset 3 (D3; *Escherichia coli*), with 4,350 genomes (http://enve-omics.ce.gatech.edu/data/public_fastani/D3.tar.gz); (iv) Dataset 4 (D4; *Bacillus anthracis*), with 468 genomes (http://enve-omics.ce.gatech.edu/data/public_fastani/D4.tar.gz); (v) Dataset 5 (D5; Parks *et al* MAGs), with 7,901 metagenome-assembled genomes (MAGs; http://enve-omics.ce.gatech.edu/data/public_fastani/D5.tar.gz).

For each dataset D1-D5, the following (meta)genomes were used as query genomes for average nucleotide identity (ANI) calculations (per the FastANI validation and benchmarking study) (Jain *et al.*, 2018): (i) Escherichia_coli_str__K_12_substr__MG1655.LargeContigs.fna for D1 (note that "GCF_000005845.2_ASM584v2_genomic.fna" was listed as the query genome in the D1 README; because there was no genome with that name in the D1 dataset, each genome in the D1 dataset was queried for "NC_000913.3", the NCBI Nucleotide accession associated with NCBI Assembly accession GCF_000005845.2. Escherichia_coli_str__K_12_substr__MG1655.LargeContigs.fna contained a chromosome with NCBI Nucleotide accession NC_000913.3, and was thus deemed to be the D1 query genome used in the FastANI study); (ii) Bacillus_anthracis_52_G_NZ_CM002395.LargeContigs.fna for D2 (per the D2 README); (iii) Escherichia_coli_0_1288_GCA_000303255.LargeContigs.fna for D3 (per the D3 README); (iv) 2000031001.LargeContigs.fna for D4 (per the D4 README); (v) Parks8k_GCA_002429585_1.LargeContigs.fna for D5 (no README was provided with D5; each MAG in the D5 dataset was therefore queried for "UBA6007", the strain name reported in Table 1 of the FastANI manuscript. Parks8k_GCA_002429585_1.LargeContigs.fna was the only MAG that contained strain name "UBA6007" and was thus deemed to be the D5 query MAG used in the FastANI study).

**Validation and benchmarking of ANI methods.** Each of the following methods was used to calculate ANI values between each (meta)genome in dataset D1-D5 and its respective query genome (see section "Acquisition of [meta]genomic data for benchmarking" above): (i) OrthoANI (OAT_cmd.jar v1.40) (Lee *et al.*, 2016); (ii) FastANI v1.33 (Jain *et al.*, 2018); (iii) skani v0.1.4 (Shaw and Yu, 2023); (iv) PyOrthoANI v0.6.0 (developed here); (v) PyFastANI v0.6.0 (developed here); (vi) Pyskani v0.1.2 (developed here). Five methods (all but Pyskani) were evaluated using 1, 8, and 16 CPUs in triplicate; Pyskani was evaluated using 1 CPU in triplicate, as skani, and thus, Pyskani, does not parallelize when performing a single pairwise distance computation (per the skani source code and as demonstrated here, Supplementary Figure S1; *n* = 717,651 total ANI computations). For each computation, "trace" in Nextflow v24.04.2 was used to log speed and memory usage (Di Tommaso *et al.*, 2017).

For (i) OrthoANI (Lee *et al.*, 2016), Java v17.0.6 was used to invoke OrthoANI from the command line, using the OAT_cmd.jar file; "-num_threads" set to 1, 8, or 16; the dataset's query genome supplied to "-fasta1"; each genome in the dataset supplied to "-fasta2"; the path to the BLAST+ v2.2.31+ executable (i.e., the BLAST+ version recommended in the OrthoANI v1.40 manual) supplied to "-blastplus_dir" (Camacho *et al.*, 2009). Pseudocode for OrthoANI is as follows:
```
java -jar OAT_cmd.jar -num_threads ${task.cpus} -fasta1 query.fna -fasta2 ref.fna
-blastplus_dir ncbi-blast/bin/ >> ref.log
```

For (ii) FastANI and (iii) skani, Singularity containers were downloaded from Bioconda (Grüning *et al.*, 2018) (fastani_1.34--h4dfc31f_0.sif and skani_0.2.0--h4ac6f70_0.sif, respectively, corresponding to the latest versions available in Bioconda at the time; note that versions reported by FastANI and skani's "--version" commands were v1.33 and 0.1.4, respectively). Both FastANI and skani were executed from the command line, with "-t" set to 1, 8, or 16; the dataset's query genome supplied to "-q"; each genome in the dataset supplied to "-r". Pseudocode for FastANI is:
```
fastANI -t ${task.cpus} -q query.fna -r ref.fna -o ref.log
```
Pseudocode for skani is:
```
skani dist -t ${task.cpus} -q query.fna -r ref.fna -o ref.log
```

For (iv) PyOrthoANI, a Singularity container containing PyOrthoANI v0.6.0, BioPython v1.84 (Cock *et al.*, 2009), and BLAST+ v2.12.0+ was constructed. PyOrthoANI was invoked directly from the command line, with "-j" set to 1, 8, or 16; the dataset's query genome supplied to "-q"; each genome in the dataset supplied to "-r". Pseudocode for PyOrthoANI is as follows:
```
orthoani -q query.fna -r ref.fna -j ${task.cpus} >> ref.log
```

Because they could not be invoked directly from the command line, (v) PyFastANI and (vi) Pyskani were each executed within a Python script. Briefly, two separate Singularity containers were constructed, each of which contained BioPython v1.84 and either PyFastANI v0.6.0 or Pyskani v0.1.2. PyFastANI was executed within a Python v3.10.12 script, where ${task.cpus} = 1, 8, or 16, as follows:
```
import sys
```

```
import pyfastani
import Bio.SeqIO
sketch = pyfastani.Sketch()
ref = list(Bio.SeqIO.parse(ref.fna, "fasta"))
sketch.add_draft("ref", (bytes(record.seq) for record in ref))
mapper = sketch.index()
query = Bio.SeqIO.parse(query.fna, "fasta")
hits = mapper.query_draft((bytes(record.seq) for record in query), ${task.cpus})
with open(ref.log, "a") as outfile:
    for hit in hits:
        print("\t".join([str(hit.name).strip(), str(hit.identity).strip(), str(hit.matches).strip(), str(hit.fragments).strip()]), file = outfile)
```

Pyskani was executed similarly (note that Pyskani, like skani, does not parallelize when performing a single pairwise distance computation; thus, only 1 CPU was used):

```
import sys
import pyskani
import Bio.SeqIO
database = pyskani.Database()
records = Bio.SeqIO.parse(ref.fna, "fasta")
ref = (bytes(record.seq) for record in records)
database.sketch("ref", *ref)
query = Bio.SeqIO.parse(query.fna, "fasta")
query_contigs = (bytes(record.seq) for record in query)
hits = database.query("query", *query_contigs)
with open(ref.log, "a") as outfile:
    for hit in hits:
        print("\t".join([str(hit.identity).strip(), str(hit.query_fraction).strip(), str(hit.reference_fraction).strip()]), file = outfile)
```

All NextFlow and Python pseudocode is available at https://github.com/lmc297/python_ani_tools.

**Aggregation and visualization of validation/benchmarking results.** Speed and memory usage results were acquired from “trace” files produced by Nextflow for each dataset/method/CPU/replicate combination (see section “Validation and benchmarking of ANI methods” above). Results were plotted in R v4.4.0 (R Core Team, 2024), using the following packages: ggplot2 v3.5.1 (Wickham, 2016), ggdist v3.3.2 (Kay, 2024), stringr v1.5.1 (Wickham, 2023a), khroma v1.14.0 (Frerebeau, 2024), gridExtra v2.3

(Auguie, 2017), forcats v1.0.0 (Wickham, 2023b). Code used to construct figures is available at https://github.com/lmc297/python_ani_tools.
