## Supplementary Figure S1 for "PyOrthoANI, PyFastANI, and Pyskani: a suite of Python libraries for computation of average nucleotide identity"

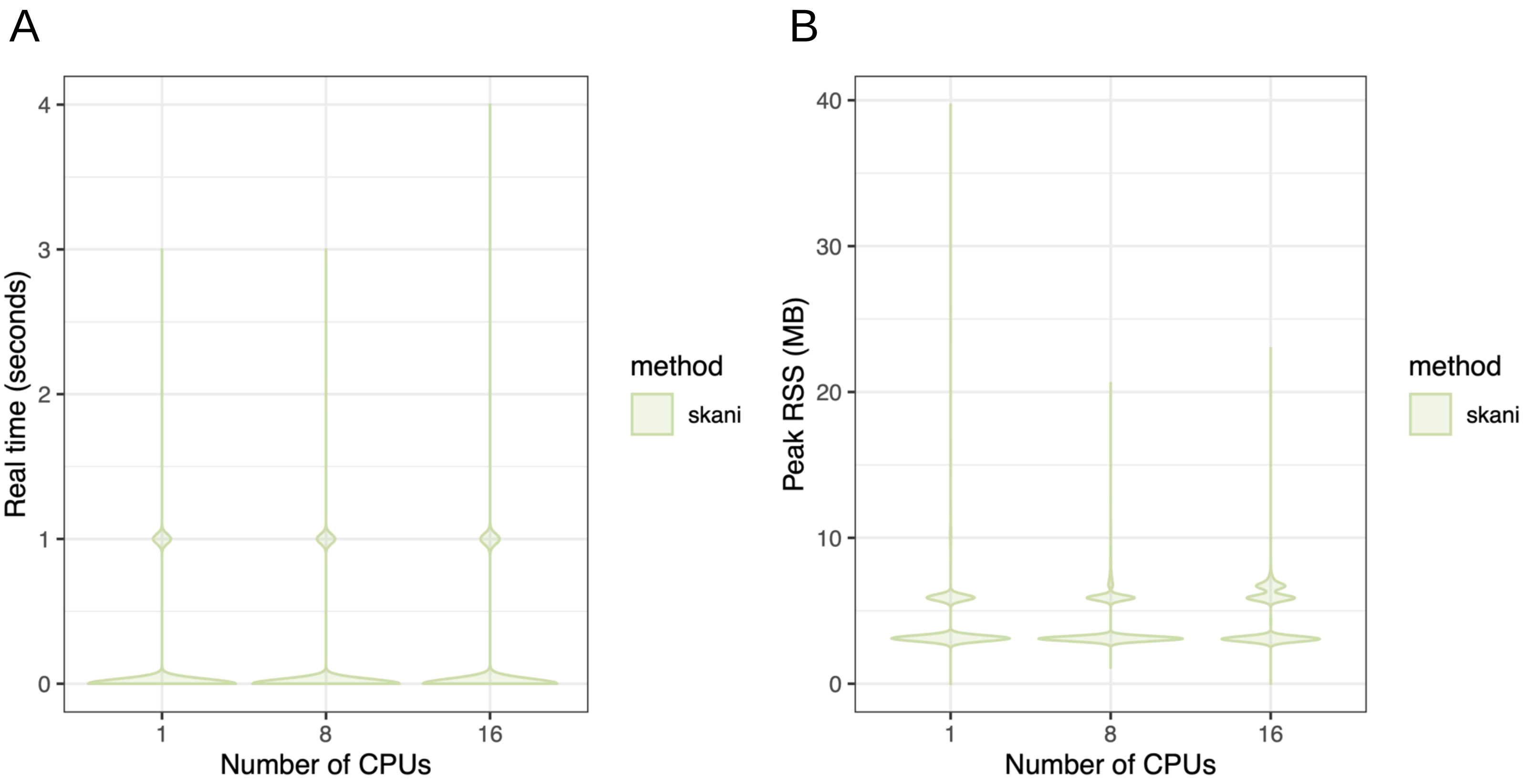


**Supplementary Figure S1.** Violin plots showing per-genome (A) time (real/wall clock time, in seconds) and (B) memory (peak resident set size [RSS], in megabytes [MB]; Y-axes) used by skani v0.1.4 on the FastANI validation/benchmarking data sets. For each genome, 1, 8, or 16 CPUs (X-axes) were specified. This analysis was performed to confirm that skani (and thus, Pyskani) does not parallelize when performing a single pairwise distance computation (per the skani source code and as demonstrated here).
