## Supplementary Figure S3 for "PyOrthoANI, PyFastANI, and Pyskani: a suite of Python libraries for computation of average nucleotide identity"

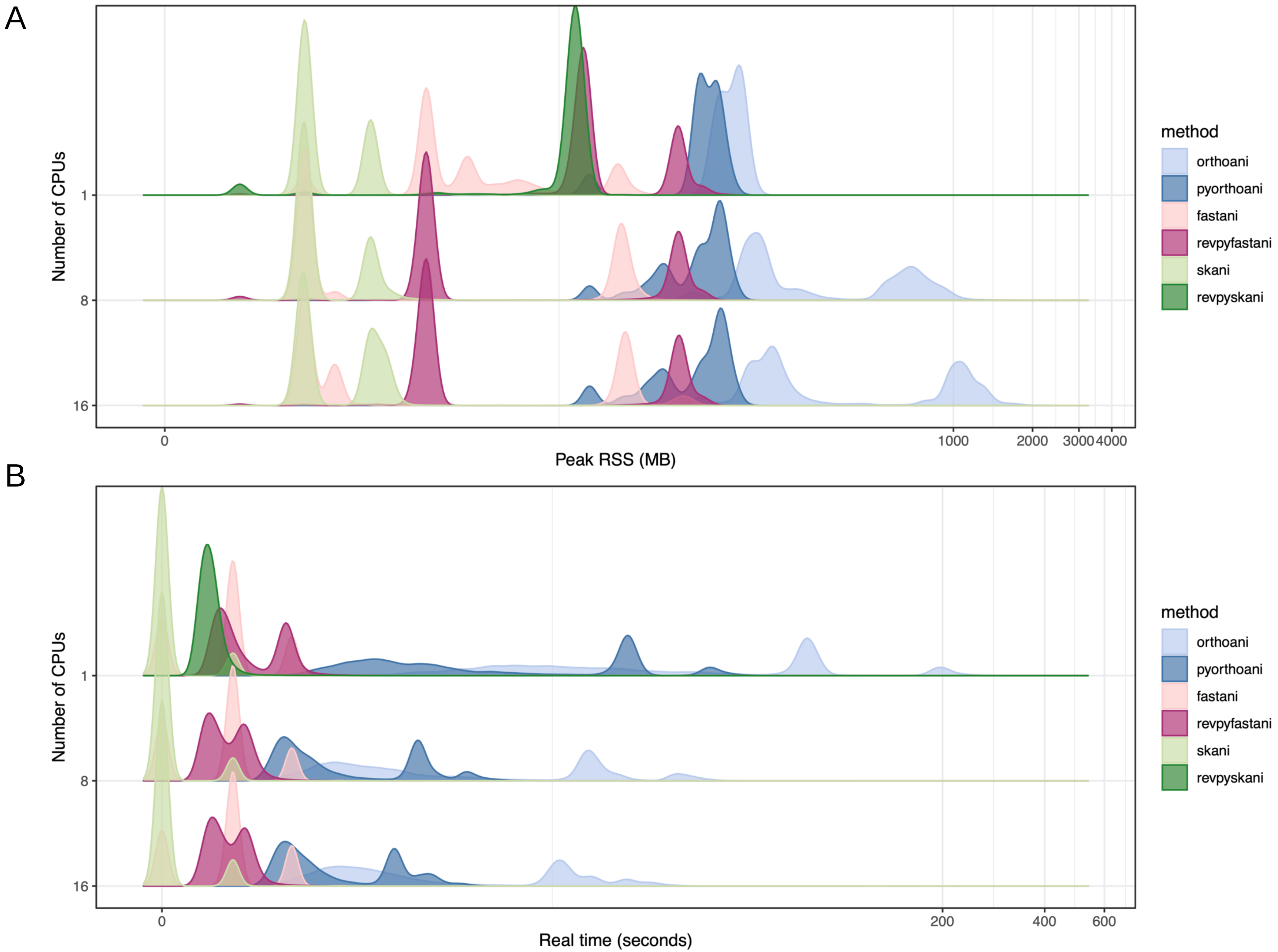


**Supplementary Figure S3.** Ridgeline plots showing per-genome (A) memory (peak resident set size [RSS], in megabytes [MB]) and (B) time (real/wall clock time, in seconds; X-axes) used by OrthoANI (orthoani), PyOrthoANI (pyorthoani), FastANI (fastani), PyFastANI (revpyfastani), skani (skani), and Pyskani (revpyskani) on the FastANI validation/benchmarking data sets. For each genome, 1, 8, or 16 CPUs (Y-axes) were specified. For Pyskani, only 1 CPU was tested, as Pyskani and skani do not parallelize when performing a single pairwise distance computation (per the skani source code and as demonstrated here).
