## Supplementary Figure S5 for "PyOrthoANI, PyFastANI, and Pyskani: a suite of Python libraries for computation of average nucleotide identity"

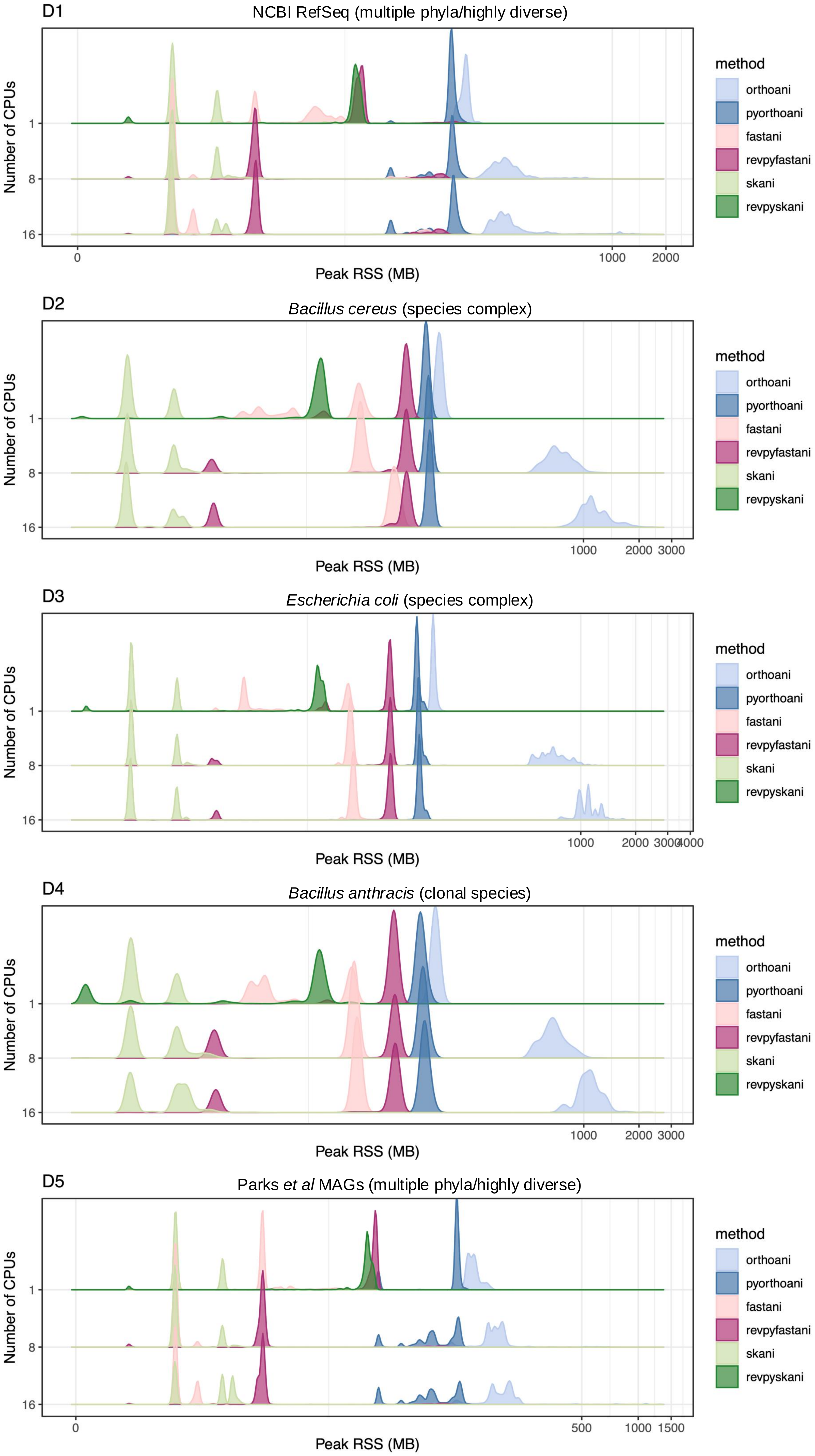


**Supplementary Figure S5.** Ridgeline plots showing per-genome memory (peak resident set size [RSS], in megabytes [MB]; X-axes) used by OrthoANI (orthoani), PyOrthoANI (pyorthoani), FastANI (fastani), PyFastANI (revpyfastani), skani (skani), and Pyskani (revpyskani) on each FastANI validation/benchmarking data set (D1-D5). For each genome, 1, 8, or 16 CPUs (Y-axes) were specified. For Pyskani, only 1 CPU was tested, as Pyskani and skani do not parallelize when performing a single pairwise distance computation (per the skani source code and as demonstrated here). Datasets: (i) D1, NCBI RefSeq (a highly diverse dataset consisting of genomes representing multiple phyla; ANI is thus not necessarily appropriate for many pairwise comparisons in this dataset); (ii) D2, *Bacillus cereus* (a species complex); (iii) D3, *Escherichia coli* (a species complex); (iv) D4, *Bacillus anthracis* (a clonal species); (v) D5, Parks *et al* MAGs (a highly diverse dataset consisting of genomes representing multiple phyla; ANI is thus not necessarily appropriate for many pairwise comparisons in this dataset).
